# A prediction model of working memory across health and psychiatric disease using whole-brain functional connectivity

**DOI:** 10.1101/222281

**Authors:** Masahiro Yamashita, Yujiro Yoshihara, Ryuichiro Hashimoto, Noriaki Yahata, Naho Ichikawa, Yuki Sakai, Takashi Yamada, Noriko Matsukawa, Go Okada, Saori C. Tanaka, Kiyoto Kasai, Nobumasa Kato, Yasumasma Okamoto, Ben Seymour, Hidehiko Takahashi, Mitsuo Kawato, Hiroshi Imamizu

**Affiliations:** Brain Information Communication Research Laboratory Group, Advanced Telecommunications Research Institute International, Kyoto, Japan.; Department of Psychiatry, Kyoto University Graduate School of Medicine, Kyoto, Japan.; Medical Institute of Developmental Disabilities Research, Showa University, Tokyo, Japan.; Department of Youth Mental Health, Graduate School of Medicine, The University of Tokyo, Tokyo, Japan.; Molecular Imaging Center, National Institute of Radiological Sciences, Chiba, Japan.; Department of Psychiatry and Neurosciences, Hiroshima University Graduate School of Biomedical & Health Sciences, Hiroshima, Japan.; Department ofPsychiatry, Graduate School of Medical Science, Kyoto Prefectural University of Medicine, Kyoto, Japan.; Department of Neuropsychiatry, Graduate School of Medicine, The University of Tokyo, Tokyo, Japan.; Computational and Biological Learning Laboratory, Department of Engineering,University of Cambridge, Cambridge, UK.; Center for Information and Neural Networks, National Institute of Information and Communications Technology Osaka, Japan.; Department of Psychology, The University of Tokyo, Tokyo, Japan

## Abstract

Individual differences in cognitive function have been shown to correlate with brain-wide functional connectivity, suggesting a common foundation relating connectivity to cognitive function across healthy populations. However, it remains unknown whether this relationship is preserved in cognitive deficits seen in a range of psychiatric disorders. Using machine learning methods, we built a prediction model of working memory function from whole-brain functional connectivity among a healthy population (*N* = 17, age 19-24 years). We applied this normative model to a series of independently collected resting state functional connectivity datasets (*N* = 968), involving multiple psychiatric diagnoses, sites, ages (18-65 years), and ethnicities. We found that predicted working memory ability was correlated with actually measured working memory performance in both schizophrenia patients (partial correlation, *ρ* = 0.25, *P* = 0.033, *N* = 58) and a healthy population (partial correlation, *ρ* = 0.11, *P* = 0.0072, *N* = 474). Moreover, the model predicted diagnosis-specific severity of working memory impairments in schizophrenia (*N* = 58, with 60 controls), major depressive disorder (*N* = 77, with 63 controls), obsessive-compulsive disorder (*N* = 46, with 50 controls), and autism spectrum disorder (*N* = 69, with 71 controls) with effect sizes *g* = −0.68, −0.29, −0.19, and 0.09, respectively. According to the model, each diagnosis’s working memory impairment resulted from the accumulation of distinct functional connectivity differences that characterizes each diagnosis, including both diagnosis-specific and diagnosis-invariant functional connectivity differences. Severe working memory impairment in schizophrenia was related not only with fronto-parietal, but also widespread network changes. Autism spectrum disorder showed greater negative connectivity that related to improved working memory function, suggesting that some non-normative functional connections can be behaviorally advantageous. Our results suggest that the relationship between brain connectivity and working memory function in healthy populations can be generalized across multiple psychiatric diagnoses. This approach may shed new light on behavioral variances in psychiatric disease and suggests that whole-brain functional connectivity can provide an individual quantitative behavioral profile in a range of psychiatric disorders.

## Introduction

Functional connectivity (FC) quantifies how brain regions are temporally coordinated, and offers potential insight into individual differences in behavior (Seeley *et al.,* 2007; Lewis *et al.,* 2009; Baldassarre *et al.,* 2012). For instance, whole-brain FC models have recently demonstrated that sets of functional connections across widespread brain regions can predict performance on cognitive tasks (Finn *et al.,* 2015; Smith *et al.,* 2015; Yamashita *et al.,* 2015; Rosenberg *et al.,* 2016). These findings suggest that specific cognitive processes may be represented by interaction patterns among distributed brain networks, and also that a general relationship exists between brain-wide connectivity and cognitive function, at least among healthy populations.

Working memory is widely accepted as one of the most important cognitive functions in daily life. Working memory reflects a brain system that temporarily maintains and processes information to guide a range of cognitive tasks (Baddeley,2003; Cowan, 2014). Increasing evidence suggest that working memory emerges from widespread brain regions that engage in sensory perception, executive control, and motor action (Owen *et al.,* 2005; Postle, 2006; Rottschy *et al.,* 2012; Nee *et al.,* 2013; D’Esposito and Postle, 2015; Eriksson *et al.,* 2015). Working memory deficits have been commonly observed in a range of psychiatric disorders (Forbes *et al.,* 2009; Millan *et al.,* 2012; Snyder, 2014; Lever *et al.,* 2015; Snyder *et al.,* 2015), and while the severity depends on the specific psychiatric diagnosis (Forbes *et al.,* 2009; Millan *et al.,* 2012; Snyder, 2014; Lever *et al.,* 2015; Snyder *et al.,* 2015), many different case-control studies have revealed dysfunction in executive control systems such as frontoparietal networks (Koshino *et al.,* 2005; Barch and Csernansky, 2007; Vasic *et al.,* 2009; De Vries *et al.,* 2014).

Functional connectivity has emerged as a promising tool in understanding the biological basis of psychiatric diagnoses, and different diagnoses have been shown to relate to unique patterns of FC (Harrison *et al.,* 2009; Baker *et al.,* 2014; Kaiser *et al.,* 2015; Yahata *et al.,* 2016). For example, a whole-brain FC-based model has been shown to predict autism spectrum disorder (ASD) (Yahata *et al.,* 2016), and specific connections within this model can predict individual clinical scores, suggesting that FC disruption is quantitatively relevant to patients behavioral abnormality. More broadly, therefore, this suggests that, a specific relationship between FC and behavior might exist across many disparate diagnoses.

With the above issues in mind, we set out to examine three mutually exclusive hypotheses about the relationship between functional connectivity models of working memory ability (FC-WMA) and observed working memory performance across healthy populations and a range of psychiatric diagnoses. The first hypothesis proposes a distinct FC-WMA relationship for each diagnosis, rationalized by the fact that each psychiatric diagnosis is characterized by differential alterations in FC (Harrison *et al.,* 2009; Baker *et al.,* 2014; Kaiser *et al.,* 2015; Yahata *et al.,* 2016) (*distinct FC*-*WMA hypothesis*). This hypothesis predicts that the FC-WMA relationship among healthy populations will fail to generalize in predicting impairments across different diagnoses. The second hypothesis proposes a common FC-WMA relationship across health and multiple diagnoses, rationalized by the fact that disruptions in fronto-parietal network-related connections are sufficient to explain the different severity of working memory deficits of different diagnoses, because fronto-parietal network dysfunction is commonly associated with working memory impairments across multiple diseases (Koshino *et al.,* 2005; Barch and Csernansky, 2007; Vasic *et al.,* 2009; De Vries *et al.,* 2014) (*common fronto-parietal network hypothesis*). This hypothesis predicts that a FC-WMA relationship, which is estimated purely from fronto-parietal network-related connections in healthy populations will generalize to predict working memory impairment across diagnoses. The third hypothesis proposes that both a common FC-WMA relationship across healthy and multiple diagnoses, *and* multiple abnormal connections among distributed networks are necessary to explain the full range of working memory deficit severities (*common whole-brain hypothesis*). This hypothesis builds on the findings that the coordinated interplay across entire brain networks underlie working memory processes (Owen *et al.,* 2005; Postle, 2006; Rottschy *et al.,* 2012; Nee *et al.,* 2013; D’Esposito and Postle, 2015; Eriksson *et al.,* 2015; Yamashita *et al.,* 2015). This hypothesis predicts that a FC-WMA relationship estimated from whole-brain functional connections in healthy populations will generalize to predict working memory impairment across diagnoses. Here, working memory impairment should be explained as an accumulation of bits of FC abnormalities, some of which are shared (e.g., fronto-parietal network-related connections), and others distinct among different diagnoses.

To test these hypotheses, we built a prediction model of working memory ability (WMA) using whole-brain FC among a healthy population, and examined its generalizability to a series of independent cohorts, with a broad variety of imaging sites, ages, ethnicities, and even psychiatric diagnoses. As we present below, the results support the third hypothesis, the *common whole*-*brain hypothesis*. We show that a whole-brain FC-WMA model is applicable across healthy populations and a range of psychiatric disorders. Moreover, detailed examination of the FC-WMA relationship across specific individual connections suggest that the working memory deficits result from combination of both diagnosis-specific and diagnosis-invariant changes in FC.

## Methods

### Design

We developed a data-driven prediction model of WMA among healthy individuals recruited at ATR (Advanced Telecommunications Research Institute International), Japan (ATR dataset; **Fig. 1A**). Independently collected resting state fMRI (rs-fMRI) was entered into this model to predict individual WMA in both healthy and clinical populations (**Fig. 1B**). Specifically, we applied the model to two independent test datasets (**Fig. 1C**): (i) healthy individuals in the USA (the Human Connectome Project dataset, HCP dataset), (ii) patients with schizophrenia, and the predicted WMA was compared with actually measured WMA. Moreover, the model was applied to patients with psychiatric diagnoses, including schizophrenia, major depressive disorder (MDD), obsessive-compulsive disorder (OCD), and ASD and also their age- and gender-matched healthy/typically developed controls (multiple psychiatric diagnoses dataset, **Fig. 1D**). The predicted WMA scores were compared with previous meta-analyses of working memory across the four psychiatric diagnoses. Throughout these three datasets, all participants provided written informed consent.

**Figure 1.**
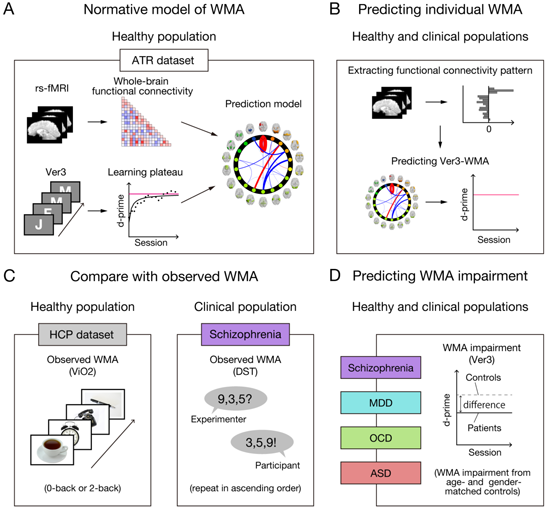
Schematic diagram of model development and application to independent datasets. (**A**) Model was developed using a whole-brain resting state FC and a learning plateau of a verbal 3-back task (Ver3-WMA) within healthy individuals from ATR dataset. (**B**) We applied the model to resting state FC patterns and predicted individual participant’s Ver3-WMA scores. (**C**) Predicted individual Ver3-WMAs were compared with observed WMAs in both healthy and clinical populations. We first examined the external validity using an independent USA dataset (HCP dataset) that included a visual-object N-back task with the performances of 0-back and 2-back conditions (ViO2-WMA). Then we examined the generalizability to a clinical population using a schizophrenia dataset that included Digit Sequencing Tests (DST-WMA). (**D**) Using the multiple psychiatric diagnoses dataset, degree of WMA impairment for each diagnosis was predicted as differences from corresponding controls. Note that the HCP dataset’s task stimuli images are just illustration purpose and different from the original stimuli.

### ATR Dataset

The first dataset was used to train the normative prediction model (*N* = 17, age 19-24 years old, 11 males).

### Behavioral assessment of working memory

The participants performed a letter-based verbal 3-back task (Ver3, **Fig. 1A**) over 25 sessions (about 80-90 min of training period in total). We evaluated their performance by calculating the d-prime for each session, and then obtained an individual learning curve by calculating 5-session moving performance averages. The individual learning curve was fitted by an inverse curve (*y* = *a* − *b*/*x*), where *y* is a d-prime in the *x*-th session, while *a* and *b* is a parameter for learning plateau and learning speed, respectively. We used the estimated learning plateau (*a*) for a measure of individual WMA. The learning plateaus showed large inter-individual differences, ranging from 1.38 to 3.87 (mean ± SD = 2.80 ± 0.72). More detailed information is described in our previous paper (Yamashita *et al.,* 2015).

### Resting state-fMRI data and Functional Connectivity (FC) estimation

We recorded a rs-fMRI scan for each participant (5 min 4 sec). All fMRI data were processed using SPM8 and in-house MATLAB code. After removing the first two volumes, the data were preprocessed with slice timing correction, realignment, and spatial smoothing with an isotropic Gaussian kernel (full width at half maximum = 8 mm). To remove several sources of spurious variance, we regressed out six motion parameters and the averaged signals over gray matter, white matter, and cerebrospinal fluid (Fox *et al.,* 2005). The gray matter signal regression improves FC estimation by effectively removing motion-related artifacts (Power *et al.,* 2014; Burgess *et al.,* 2016; Ciric *et al.,* 2017). Finally, we performed “scrubbing” (Power *et al.,* 2012) in which we removed scans where framewise displacement was > 0.5 mm.

Based on the 18 whole-brain intrinsic networks of BrainMap ICA (Laird *et al.,* 2011), we calculated FC values for pair-wise between-network (18 × 17/2 = 153) connections and within-network connections. Between-network FC was calculated as Pearson’s correlation between blood-oxygen-level dependent signal time courses averaged across voxels within each network, and then transformed to Fisher *Z* values. Within-network FC was calculated as mean voxel-wise correlations within each network as follows. First, the time series for a voxel was correlated with every other voxel within a network, and this calculation was repeated for every voxel in the network. Next, the correlation coefficients were transformed into Fisher *Z* values. By averaging the *Z* values within each network, within-network FC for all 18 networks was calculated.

### Developing prediction model

To predict individual learning plateaus in the Ver3-WMA, we performed a sparse linear regression analysis (Sato, 2001) (VBSR toolbox; http://www.cns.atr.jp/cbi/sparseestimation/sato/VBSR.html) on the whole-brain FC values. In this framework, individual Ver3-WMAs were modeled as a linear weighted summation of a small number of FC values among the intrinsic networks. To build a single prediction model, we utilized all the data (*N* = 17) as the training set. More detailed information on the model development is described in our previous paper (Yamashita *et al.,* 2015).

### Human Connectome Project (HCP) Dataset

The second dataset was collected in the HCP (Van Essen *et al.,* 2013), which is publicly available as “500 Subjects Release” (*N* = 542). We restricted our analysis to participants for whom all rs-fMRI, visual-object N-back (HCP: WM_Acc), and Raven’s progressive matrices with 24 items (HCP: PMAT24_A_CR) (Bilker *et al.,* 2012) were available (*N* = 474; 194 males, 5-year age ranges in the Open Access Data: 22-25, 26-30, 31-35 and 36+ years old).

### Behavioral assessment of working memory

Individual WMA was briefly measured by the visual-object N-back with 0-back and 2-back conditions (ViO2-WMA), which consisted of four different categorical stimuli: faces, body parts, tools, or places (**Fig. 1C**). The N-back task was performed in two runs of fMRI experiment, with 5 min scan acquisition for each run. The scores were evaluated by the accuracy percentage of 2-back and 0-back conditions (86.0% ± 9.5% (SD), range 45.8% to 100%) and showed non-normal distributions. Thus, we performed nonparametric statistical test, Spearman’s rank correlation, to examine the correlation of the model predictions with ViO2-WMA scores. Additionally, general fluid intelligence was assessed by Raven’s progressive matrices. The scores are integers that indicate the number of correct items (16.5 ± 4.8 (SD) from 4 to 24).

### Resting state-fMRI data and Functional Connectivity (FC) estimation

We used rs-fMRI data that were already preprocessed and denoised by the FIX procedure (Salimi-Khorshidi *et al.,* 2014). Realignment and spatial smoothing was performed with an isotropic Gaussian kernel (full width at half maximum = 4 mm). Nuisance regression was performed using average signals of gray matter, white matter, and cerebro-spinal fluid. Finally, a band-pass filter (0.009-0.08 Hz) was applied and volumes with framewise displacement > 0.5 mm were removed. FC estimation was performed in the same way as in the ATR dataset.

### Examination of model prediction

We entered whole-brain FC values into the prediction model developed from the ATR dataset, and predicted individual’s Ver3-WMA. To examine whether the model significantly predicted individually measured ViO2-WMA, we shuffled the participant labels of the model predictions and compared the predicted Spearman’s rank correlation coefficient with the obtained null distribution by 10,000 permutations.

### Multiple Psychiatric Diagnoses Dataset

We recruited participants with four different psychiatric diagnoses: schizophrenia, MDD, OCD, and ASD. These data were collected across five universities and seven scanners among a Japanese neuropsychiatry consortium (Yahata *et al.,* 2016) (**Supplementary Tables 2** for demographic data; http://www.cns.atr.jp/decnefpro/). Patients and their age- and gender-matched healthy controls were recruited at each site: 58 patients with schizophrenia (age 37.9 ± 9.3 (standard deviation, SD), 30 males) and 60 controls (age 35.2 ± 8.4, 40 males) were recruited at Kyoto University; 77 patients with MDD (age 41.6 ± 11.2, 43 males) and 63 controls (age 39.3 ± 12.0, 29 males) were recruited at Hiroshima University; 46 patients with OCD (age 32.2 ± 9.9, 17 males) and 50 controls (age 30.0 ± 7.3, 24 males) were recruited at Kyoto Prefectural University of Medicine. Patients with ASD and their controls were recruited at two sites; 33 patients with ASD (age 32.8 ± 8.4, 21 males) and 33 controls (age 34.7 ± 7.0, 18 males) were recruited at the University of Tokyo, and 36 patients with ASD (age 29.9 ± 7.2, 36 males) and 38 controls (age 32.5 ± 7.4, 38 males) were recruited at Showa University. Additional details including inclusion/exclusion criteria, medication usage, and informed consent are described in **Supplementary Methods**.

### Resting state-fMRI data and Functional Connectivity (FC) estimation

We collected rs-fMRI data for each participant (for scanning parameters see **Supplementary Table 3**). Preprocessing and FC estimation were performed in the same way as in the ATR dataset. We conducted quality control for the rs-fMRI data and excluded participants if more than 40% of their total number of volumes of their rs-fMRI data were removed by the scrubbing method. After their data quality was assured, age-and gender-matched healthy control subjects were included in the analysis.

### Behavioral assessment of working memory

In the schizophrenia patients, general cognitive ability was assessed by the Japanese version of Brief Assessment of Cognition in Schizophrenia (BACS-J) (Kaneda *et al.,* 2007). This cognitive battery is composed of six subtests including Digit Sequencing Test (DST) - a WMA measure. In the DST, sequences of numbers were auditorily presented, with increasing length from three to nine digits (**Fig. 1C**). Participants repeated the sequences aloud by sorting in ascending order. The DST-WMA scores were integers that indicate the number of correct trials among 28 trials (18.4 ± 4.1 (SD),range 10 to 27). We examined the statistical significance on the prediction accuracy by permutation tests as described in HCP dataset. Furthermore, to examine whether the model specifically predicted DST-WMA independently of age and general cognitive performance, we performed a partial correlation analysis that factored out these two variables. Here, general cognitive performances were evaluated by composite score of BACS’s five subtests other than the DST. The composite score was obtained by averaging over five *Z*-scores of the subtests.

### Examination of model prediction

In the model predictions, we detected outliers within each group (defined as values > 3 SD from the mean) for a control participant of MDD and a patient with OCD (*N* = 1, 1, respectively). These two participants were excluded from further analysis. Patients’ predicted Ver3-WMA differences were evaluated by the *Z*-scores standardized to their age-and gender-matched controls collected in the same site. After confirming the homoscedasticity (Bartlett’s test, *P* = 0.19), the standardized WMA differences were entered in a one-way ANOVA with diagnosis as a between-participant factor. Post-hoc pair-wise comparisons were corrected using Holm’s method.

## Results

### Building Prediction Model Using Whole-Brain Functional Connectivity (FC)

We used the ATR dataset as the training set to develop a normative model of individual WMA. Using sparse linear regression, individual Ver3-WMAs were modeled as a linear weighted summation of automatically selected 16 FC values among 15 intrinsic networks (**Fig. 2**). Ver3-WMA scores were positively correlated with three FC values (P1-P3) and negatively correlated with the remaining 13 FC values (N1-N13). Each network of the connections and the anatomical regions in the network are summarized in **Table 1** and **Supplementary Table 1**, respectively. For example, FC within a left fronto-parietal network (P1) accounted for about 34% of the total variance of the model predictions. We did not find a significant correlation between the predicted Ver3-WMA scores and age (*r* = 0.21, *P* = 0.42), gender (*r* = 0.28, *P* = 0.28), or head motion (*r* = − 0.37, *P* = 0.14). This provided a normative prediction model based on healthy young Japanese participants.

**Figure 2.**
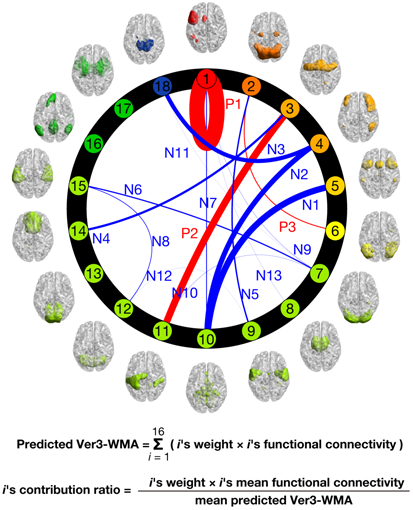
Circle plots of normative model of Ver3-WMA. Individual Ver3-WMAs are predicted by a linear weighted summation of 16 FC values. Connection thicknesses indicate contribution ratios (weight x FC at each connection: **Table 1**). Connections are labeled “Positive/Negative (P/N)” based on correlation coefficient signs with Ver3-WMA scores, whereas numbers indicate descending orders of contribution ratio magnitudes. Each network’s color indicates relevance with working memory function based on BrainMap ICA (Laird *et al.,* 2011); warmer colors indicate closer relevance to working memory function. Each network’s label and regions included in it are summarized in **Supplementary Table 1**.

### Prediction in Independent Test Set of Healthy Individuals

We next tested the model’s generalizability to an entirely independent healthy cohort (HCP dataset). We found that the model significantly predicted ViO2-WMA scores (Spearman’s rank correlation, *ρ* = 0.09, *P* = 0.030). However, we found that ViO2-WMA scores were also positively correlated with general fluid intelligence (*ρ* = 0.46, *P* = 3.3 × 10^−26^) and negatively correlated with average in-scanner head motion (*ρ* = −0.24, *P* = 1.5 × 10^−7^). To examine whether the model specifically predicted individual WMA, we performed a partial correlation analysis while factoring out these two variables. This revealed a more significant partial correlation between the predicted Ver3-WMA scores from FC and the measured ViO2-WMA scores (Spearman’s rank partial correlation *ρ* = 0.11; *P* = 0.0072, **Fig. 3A**). The null distributions created for all the permutation tests are illustrated in **Fig. 3B**. Therefore, the model captures FC variations specific to WMA independently of general fluid intelligence and head motion.

**Figure 3.**
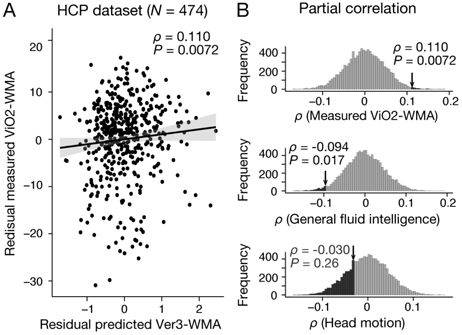
Generalizability to HCP dataset. (**A**) Significant Spearman’s rank partial correlation between predicted Ver3-WMA and measured ViO2-WMA scores while factoring out general fluid intelligence and head motion (*ρ* = 0.110, *P* = 0.0072). (**B**) Frequency of Spearman’s rank partial correlation of predicted Ver3-WMA with observed ViO2-WMA, general fluid intelligence, and head motion over 10,000 permutations. Bin width is 0.005.

### Prediction in Independent Test Set of Individual Patients with Schizophrenia

We also addressed whether the model could predict individual WMA differences among patients with a single psychiatric diagnosis, focusing on schizophrenia. We applied the model to an independent test set of schizophrenia patients in the multiple psychiatric patient dataset. We found that the model prediction was positively correlated with DST-WMA scores (*r* = 0.26, *P* = 0.025). However, DST-WMA scores also correlated positively with general cognitive ability in BACS (*r* = 0.60, *P* = 5.4 × 10^−7^), negatively with age (*r* = −0.37, *P* = 0.005), but not with head motion (*r* = −0.02, *P* = 0.90). To confirm that the model predictions were specific to WMA, we examined the relationship between predicted Ver3-WMA scores from FC, the measured DST-WMA scores, age, and general cognitive ability. After controlling the age and the general cognitive ability using a partial correlation analysis, the model predictions showed significant correlations with DST-WMA scores (*ρ* = 0.25, *P* = 0.033, **Fig. 4A**). The null distributions created for all the permutation tests are illustrated in **Fig. 4B**. Therefore, the model captures FC variations that are specific to WMA independently of age or general cognitive ability.

**Figure 4.**
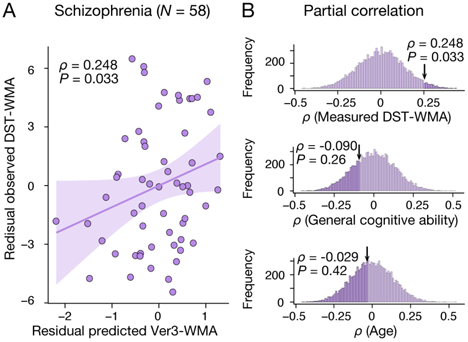
Generalizability to Schizophrenia dataset. (**A**) Significant Pearson partial correlation between predicted Ver3-WMA and measured DST-WMA scores while factoring out general cognitive ability and age (*ρ* = 0.248, *P* = 0.033). (**B**) Frequency of Pearson partial correlation of predicted Ver3-WMA with observed DST-WMA, general fluid intelligence, and head motion while factoring out two other variables over 10,000 permutations. Bin width is 0.01.

### Prediction in Independent Test Set of Four Distinct Psychiatric Disorders

We addressed whether our model could quantitatively reproduce degrees of WMA deficits across four psychiatric diagnoses. WMA impairments have generally been observed, in descending order of severity, in schizophrenia, MDD, OCD, and ASD (Forbes *et al.,* 2009; Snyder, 2014; Lever *et al.,* 2015; Snyder *et al.,* 2015). First, we compared the model predictions between patients and controls for each diagnosis scanned at the same site to remove scanner and imaging protocols as disturbance variables. We identified significant differences in the model predictions between the patient and normal controls only for schizophrenia patients (two-tailed *t*-test for schizophrenia group: *t*_116_ = −3.68, *P* = (3.5 × 10^−4^) × 4 = 0.0014, Bonferroni corrected; **Fig. 5A)**. Second, we calculated individual patients’ *Z*-score of the predicted WMAs for each diagnosis (**Fig. 5B**). A one-way ANOVA revealed a significant main effect of diagnosis on the *Z*-score (*F*_3,245_ = 7.49, *P* = 8.15 × 10^−5^). The severity of the predicted impairment in schizophrenia patients was larger than all other diagnoses (post-hoc Holm’s controlled *t*-test, adjusted *P* < 0.05).

The predicted WMA alteration was more negative in the order of schizophrenia, MDD, OCD, and ASD with effect sizes (Hedge’s g) of −0.68, −0.29, −0.19, and 0.09, respectively. Assuming that previous meta-analyses on digit span tasks (DSp-WMA) (Forbes *et al.,* 2009; Snyder, 2014; Snyder *et al.,* 2015) provide a ground-truth for WMA impairment severity, the model predictions were quantitatively supported. Specifically, the predicted Ver3-WMA alteration effect sizes fell within confidence intervals of estimates for forward DSp-WMA in schizophrenia, MDD, and OCD and for backward DSp-WMA in OCD (**Fig. 5C**). Note that we only compared these three diagnoses since no meta-analysis was available for ASD. We identified no significant differences in head motion between patients and their healthy controls in any diagnosis (two-tailed *t*-tests, *P* > 0.32, Bonferroni corrected). Importantly, the model was determined by capturing a normal range of variation in FC and WMA and reproduced an abnormal range of variation in FC and WMA, considering not only the order but also the quantitative aspects of WMA deterioration across the four distinct diagnoses.

**Figure 5.**
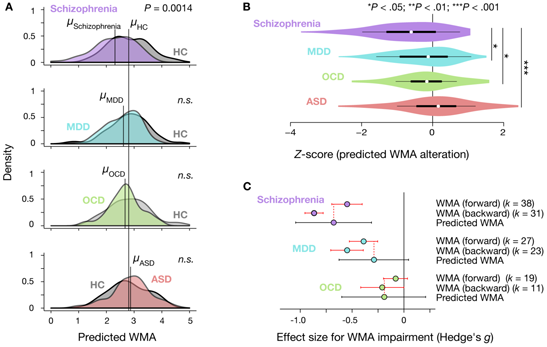
Quantitative prediction of diagnosis-specific alterations of WMA. (**A**) Predicted WMA for patients (*N* = 58, 77, 45, and 69 for schizophrenia, MDD, OCD, and ASD, respectively) and their age- and gender-matched healthy/typically developed controls (HC, *N* = 60, 62, 51, and 71) shown as kernel density. For illustration purposes, distribution of each HC was standardized to that of the ATR dataset, and the same linear transformation was applied to patients’ distributions. *μ* indicates mean value for each group. Significant impairment was predicted only in schizophrenia patients (*t*_116_ = −3.68, *P* = 0.0014, Bonferroni corrected). (**B**) Violin plots of *Z*-scores for predicted WMA alterations. We observed a significant main effect of diagnosis (one-way ANOVA, *F*_3,245_ = 7.45, *P* = 8.58 × 10^−5^) and significant differences between schizophrenia and all other diagnoses (Holm’s controlled *t*-test, *P* < 0.05). White circles indicate medians. Box limits indicate 25th and 75th percentiles. Whiskers extend 1.5 times interquartile range from 25th and 75th percentiles. (**C**) Comparison of estimated effect sizes for WMA deficits. *k* indicates number of studies included in the metaanalyses (Forbes *et al.,* 2009; Snyder, 2014; Snyder *et al.,* 2015). Error bars indicate 95% confidence intervals.

**Figure 6.**
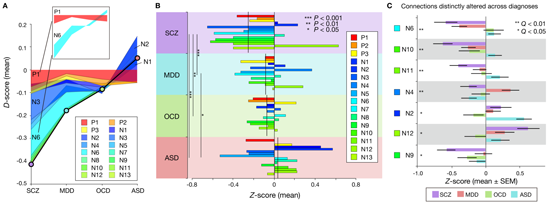
Accumulation of both shared and distinct Function Connectivity (FC) differences exhibits diagnosis-specific WMA. (**A**) Accumulation of averaged *D*-scores for all 16 connections. Bold black line indicates summation of contributions by all connections, corresponding to predicted WMA alteration. This figure shows how diagnosis-specific WMA impairment results from complex disturbances of multiple connections. Upper panel depicts two representative alteration patterns across diagnoses. While connection P1 commonly decreased WMA across diagnoses, connection N6 distinctly altered WMA (decrease in schizophrenia and MDD and increase in OCD and ASD). (**B**) *Z*-scores (normalized *D*-scores) for each diagnosis. Left asterisks and lines indicate significant differences in mean *Z*-scores between two diagnoses (*P* < 0.05, Bonferroni corrected). Vertical lines across horizontal bars indicate *Z*-scores averaged across connections. (**C**) *Z*-scores for each connection, sorted by small *P* values of diagnosis effect (Kruskal-Wallis test, *Q* < 0.05, FDR corrected).

We conducted additional control analyses in which we made different prediction models: an fronto-parietal network *model* using only fronto-parietal network-related FC values, and an fronto-parietal network-*removed model* using whole-brain FC values with fronto-parietal network-related connections removed (see **Supplementary Results**). The fronto-parietal network-removed model failed to yield significant prediction performance within the discovery cohort. The fronto-parietal network model yielded significant prediction performance among the discovery cohort, but still failed to predict WMA in individual patients with schizophrenia. Nor could the fronto-parietal network model reproduce the ordered severity of WMA impairments for the four diagnoses (see **Supplementary Results**). These results suggest that fronto-parietal network-related connections are necessary to predict individuals’ WMA but insufficient to predict WMA deficits across multiple psychiatric diagnoses.

### Characterizing WMA-Related Functional Connectivity (FC) Patterns in Psychiatric Diagnoses

Given a common FC-WMA relationship across these diagnoses, we examined how diagnosis-dependent WMA impairment resulted from FC alteration patterns that characterize each diagnosis. Since the model is the weighted summation of 16 FC values, increased/decreased WMA reflects the sum of the increased/decreased weighted-FC values of all the connections. To investigate the effect of each connection on WMA impairment, for each individual patient, we examined the difference in weighted-FC value at each connection from the corresponding control average. We call this difference the *D*-score (see Supplementary Methods). By averaging the *D*-scores within each diagnosis, **Fig. 6A** shows that diagnosis-dependent WMA impairment is attributed to the accumulation of diagnosis-dependent *D*-scores for all the connections. Some *D*-scores are relatively constant, while others are variable across diagnoses. For example, the *D*-score for P1 was commonly negative regardless of the diagnosis, while the *D*-score for N6 was negative or positive, dependent on the diagnosis (inset, **Fig.6A**). Therefore, **Fig. 6A** qualitatively suggests that diagnosis-dependent WMA impairment is derived from complex FC alterations patterns, where some connections are altered in a common manner across other diagnoses, while others are altered distinctly across diagnoses.

Furthermore, we standardized the *D*-score and entered the *Z*-score in a two-way ANOVA with diagnosis as a between-participant factor and connection as a within-participant factor (see **Supplementary Methods**). We found a significant main effect of diagnosis (*P* < 1.0 × 10^−4^; **Fig. 6B**). Schizophrenia patients showed significantly more negative mean *Z*-scores across connections than the other diagnoses (post-hoc diagnosis-pair-wise comparisons, *P* < 0.05). This suggests that the global patterns of the WMA-related 16 connections in schizophrenia were more severely disrupted than the other three diagnoses. Additionally, MDD patients showed significantly more negative mean Z-scores than ASD patients, suggesting that MDDs have more severe deterioration in general WMA-related FC compared to ASD.

Moreover, we found a significant interaction effect between diagnosis and connection in the Z-scores (*P* < 1.0 × 10^−5^; **Fig. 6C**), suggesting that FC alterations at particular connections are diagnosis-dependent. In seven connections that span 11 intrinsic networks (**Fig. 6C**), the *Z*-scores were significantly different among the four diagnoses (*Q* < 0.05, FDR corrected), suggesting that these connections are differentially altered across the diagnoses. Conversely, no significant differences in the *Z*-scores were observed across diagnoses in the remaining nine connections that span 13 networks, suggesting no significant effect of diagnosis on these FC difference *Z*-scores. For example, the positive recurrent connectivity within the left fronto-parietal network (P1), which is crucial for executive functions and is commonly dysfunctional across psychiatric diagnoses (Anticevic *et al.,* 2014; Baker *et al.,* 2014; Kaiser *et al.,* 2015), was commonly weaker than the controls and associated with a lower WMA. However, P1 deterioration accounted for just a part of the WMA impairment (7.5-34.9% of the total amount of negative *D*-scores in each diagnosis).

Illustrating FC-WMA relationships for each individual connection, Fig. 6 together indicates that the FC alterations are either shared (e.g., P1) or distinct (e.g., N6) across diagnoses. These results suggest that each FC alteration may be differently related to WMA across diagnoses. In contrast, by combining these diagnosis-invariant and diagnosis-specific patterns of FC alterations, the whole-brain model coherently transforms them into WMA alterations under the common whole-brain FC-WMA relationship.

## Discussion

We investigated the brain-behavior relationship between FC and WMA across healthy populations and four psychiatric diagnoses. We built a prediction model of WMA using data-driven analysis of whole-brain FC among healthy Japanese individuals (ATR dataset). Using rs-fMRI scans independently collected at different sites, our model predicted individual differences of WMA not only in a healthy population (HCP dataset), but also in schizophrenia patients (schizophrenia dataset). It also reproduced the order and the degrees of WMA impairment for four distinct diagnoses (multiple psychiatric diagnoses dataset). These model predictions were not explained purely by age, general intellectual/cognitive ability, or in-scanner head motion. Our results provide the first evidence for the presence of a general whole-brain FC-WMA relationship across healthy populations and a range of psychiatric disorders. That is, our results support the idea that WMA impairment in psychiatric disorders is a continuous deviation from a normal pattern while preserving the common relationship between brain-wide connectivity and WMA. Further investigation into how altered FC resulted in WMA impairment revealed overall disruptions in WMA-related connections in schizophrenia patients, suggesting that their severe WMA deficits are derived from distributed brain networks rather than local areas. Across diagnoses, altered FC at each individual connection showed a mixture of diagnosis-invariant and diagnosis-specific FC alteration patterns, suggesting that whole-brain modeling is required to coherently link such complex network disturbances with WMA.

Our results support the common whole-brain hypothesis amongst the three proposed competing hypotheses, for several reasons. Our normative model built from healthy populations generalized to predict WMA impairments across distinct psychiatric diagnoses. It predicted individual differences in WMA among schizophrenia patients as well as a diagnosis-dependent pattern of WMA impairment in four distinct diagnoses. These results do not support the hypothesis that the whole-brain FC-WMA relationship is specific to each diagnosis (*distinct hypothesis*) but support two alternative hypotheses where the relationship is common to healthy populations and multiple diagnoses. Further investigations of individual FC alterations that cause WMA impairment identified both shared and distinct patterns of abnormality across diagnoses. These altered FC spanned widely distributed brain networks rather than restricted connections related to the fronto-parietal network. A control analysis using only fronto-parietal network-related FC yielded no significant prediction of diagnosis-dependent patterns in WMA (see **Supplementary Results**). These results do not support the hypothesis that WMA is simply determined by fronto-parietal network-related connections independently of diagnosis (*common fronto*-*parietal network hypothesis*), but instead support the hypothesis that diagnosis-dependent WMAs can be explained by multiple connections among widely distributed brain networks (*common whole*-*brain hypothesis*).

Our model achieved generalization to test sets that differed from the training sets in terms of ethnicities, imaging sites, age groups, working memory tasks, and psychiatric diagnoses. Though prediction accuracy in a test set is inherently smaller than training accuracy (Gabrieli *et al.,* 2015), our model’s accuracy was comparable with a previous neuroimaging marker of attention ability for an external test set (*r* ~ 0.3) (Rosenberg *et al.,* 2016). Moreover, our model provided quantitative working memory impairments across the different diagnoses. Specifically, the effect sizes of working memory impairment predicted by our model in schizophrenia, MDD, and OCD patients were comparable with previous meta-analyses on a digit span forward condition that assesses verbal WMA (Forbes *et al.,* 2009; Snyder, 2014; Snyder *et al.,* 2015).

Although no meta-analysis was identified in ASD, previous studies generally showed few differences in verbal WMA from typically developed controls (Koshino *et al.,2005*; Williams *et al.,* 2005; Lever *et al.,* 2015), consistent with the predictions of our model. Here we propose two possible reasons for the moderate generalizability of our normative model. First, since its network definition is based on thousands of fMRI experiments (BrainMap ICA) (Laird *et al.,* 2011), FC is estimated based on general-population intrinsic functional networks and allows generalization beyond ethnicities, imaging sites, and age. Second, we spent 90 minutes measuring each participant’s WMA in the discovery dataset (Yamashita *et al.,* 2015). Such precise WMA measurements in the training data ensured that our model captured essential FC for WMA.

Larger sample sizes in the training set generally improves prediction accuracy in machine learning (Bishop, 2006). Using identical methods in the ATR dataset (*N* = 17), we tried to develop a new prediction model of ViO2-WMA using the HCP dataset as a training dataset (*N* = 474). However, this model failed to provide significant prediction within the training samples (*R*^2^ = 0.005). This seemingly unexpected result may partly result from differences in the way individuals’ WMA were evaluated. The HCP conducted the N-back task in a limited time (10 min), which may be insufficient for developing precise prediction model of individuals’ cognitive ability. In fact, with careful examination of cognitive performance, Rosenberg et al. built a model using modest (*N* = 25) training samples and demonstrated robust generalization to independent test sets (Rosenberg *et al.,* 2016).

We carefully excluded the possibility of spurious correlations (Whelan and Garavan, 2014; Siegel *et al.,* 2016). At an individual level, we examined general intellectual/cognitive ability, age, and head motion and confirmed that these disturbance variables had a minimal effect on prediction. At a group level, we analyzed age- and gender-matched healthy/typically-developed controls from the same sites and compared the alterations from the controls (*Z*-scores), thereby minimizing the false positives that could be derived from age, gender, or imaging sites/parameters.

Schizophrenia is characterized by a more substantial impairment in working memory than other psychiatric disorders (Forbes *et al.,* 2009; Millan *et al.,* 2012; Park and Gooding, 2014), in keeping with the notion that this impairment may be an endophenotype in schizophrenia (Park and Gooding, 2014). Our model predicted individual differences in WMA in this diagnosis, suggesting that the abnormal range of WMA variations is continuous with WMA variations within normal populations. Note that our model of an individual WMA predicted significant group-level impairment, providing a direct link between individual brain connectivity and the characteristic symptoms of schizophrenia. The magnitude of the disruption in the network connections was generally more severe in schizophrenia than in any other diagnoses. Our results provide unique evidence that WMA impairment in schizophrenia is an extreme deviation from a normal pattern of the FC-WMA relationship.

A single behavioral abnormality can result from a complex pattern of multiple FC alterations. We showed that the diagnosis-dependent pattern of WMA impairment results from the mixture of diagnosis-specific and diagnosis-independent differences in FC. Specifically, distinct alterations were identified in seven connections (**Fig. 6C**), suggesting that characteristic FC alterations in each diagnosis contribute to different degrees of WMA impairment. This explains why distinct diagnoses show a diagnosis-dependent level of WMA impairment. Conversely, no significant FC differences were identified in the remaining nine connections across diagnoses. Since most of these were altered toward lower WMAs (i.e., negative *Z*-scores), they generally contributed to WMA impairment, explaining why a range of psychiatric disorders commonly shows WMA impairment. Importantly, such neurobiological insights into behavioral abnormality are consistent with recent transdiagnostic studies of genomics (Plomin *et al.,* 2009; Smoller *et al.,* 2013; O’Donovan and Owen, 2016) and neuroimaging (Goodkind *et al.,* 2015; Clementz *et al.,* 2016), which indicate that some neurobiological changes are shared across psychiatric diagnoses and that the behavioral abnormality in these diseases are quantitative traits rather than qualitative conditions.

Our proposed two-stage approach, which builds a normative model and applies it to multiple diagnoses, is an effective technique to systematically compare neural substrates across multiple diagnoses under a unified framework. Clinical measures of attention deficit hyperactivity disorder were previously predicted by FC patterns that determine attention ability in healthy populations (Rosenberg *et al.,* 2016), suggesting common FC-cognition relationships across healthy and clinical populations. However, that study failed to directly examine the attention ability for participants with attention deficit hyperactivity disorder and was restricted to a single diagnosis. We investigated individual WMA in schizophrenia patients and examined it across multiple distinct psychiatric diagnoses. By combining whole-brain intrinsic FC patterns, machine-learning techniques, and relatively large independent samples including multiple psychiatric diagnoses (*N* = 969), our predictive modeling provides a general mechanistic explanation about cognitive abilities across healthy populations and psychiatric diagnoses. In this framework, cognitive deficits in a range of psychiatric disorders should be recognized as a relative quantitative deviation from normal patterns of FC-cognitive ability relationships. By coherently establishing FC-cognition relationships from normal to abnormal, our two-stage approach could potentially cluster multiple psychiatric disorders based on neurobiological measures and behaviors (Insel *et al.,* 2010; Insel and Cuthbert, 2015). Note that since network identification was performed on healthy participants, there may be other diagnostic-specific intrinsic network connections that are blind to the testing phase of this approach. These untested issues could be better understood in the future by examining diagnosis-specific model’s generalizability to other diagnosis or to healthy populations.

In our study, site differences could produce two types of confound: 1) from measurement settings such as scanners or protocols, and 2) population sampling bias such as participant recruitment. We analyzed the standardized difference (*Z*-score) between patients and healthy controls that were examined in the same site to eliminate site differences. This analysis is effective for the first type of confound because this confound is common to patients and controls. Therefore, our results cannot be explained by the differences in scanner type. However, our analysis cannot eliminate the second instance of confound when FC values in each control group are not identically distributed across different sites due to the sampling bias at each site. This issue is unresolved in our current study.

In conclusion, our data provide a unified WMA framework across healthy populations and multiple psychiatric disorders. Our whole-brain FC model quantitatively predicted individual WMA in independently collected cohorts of healthy populations and patients with any of four psychiatric diagnoses (*N* = 969). Our results suggest that the typical FC-WMA relationship identified in healthy populations is commonly preserved in these psychiatric diagnoses and that WMA impairment in a range of psychiatric disorders can be explained by the cumulative effect of multiple disturbances in FC among distributed brain networks. Our findings lay the groundwork for future research to develop a quantitative, brain-wide-connectivity-based prediction model of human cognition that spans health and psychiatric disease.

## Acknowledgements

We thank the Human Connectome Project, WU-Minn Consortium (principal investigators, D. Van Essen and K. Ugurbil; 1U54MH091657; http://www.humanconnectome.org/) for providing us the access to the data.

